# Generation of Infectious Prions Amenable to Site-specific Click Chemistry

**DOI:** 10.64898/2026.04.08.717205

**Authors:** Ryan G. Campbell, Jolene N. Iseler, Abigail M. Schwind, Surachai Supattapone

## Abstract

Prion diseases are a group of fatal neurodegenerative diseases that proceed through the templated conversion of the normal PrP^C^ protein to a self-propagating and infectious form, termed PrP^Sc^. This conversion process is central to disease progression. However, due to difficulties in producing functional PrP^Sc^ molecules that can be selectively modified with chemical probes, many aspects of PrP^Sc^ biology cannot be directly studied. To overcome this limitation, we substituted p-azido-L-phenylalanine (AzF), a small click chemistry-reactive amino acid, for tryptophan residue 99 of PrP^C^. W99AzF PrP^C^ substrate can efficiently and faithfully propagate either infectious or non-infectious PrP^Sc^ conformers *in vitro*. Critically, W99AzF PrP^Sc^ amyloid fibrils remain amenable to click chemistry by various ligands after the prion conversion process. Through the combination of site-specific substitution, the modularity of click chemistry, and the functional diversity of click labels, a multitude of modified prions can now be produced to ask targeted questions about the biochemical and biological basis of prion infectivity.

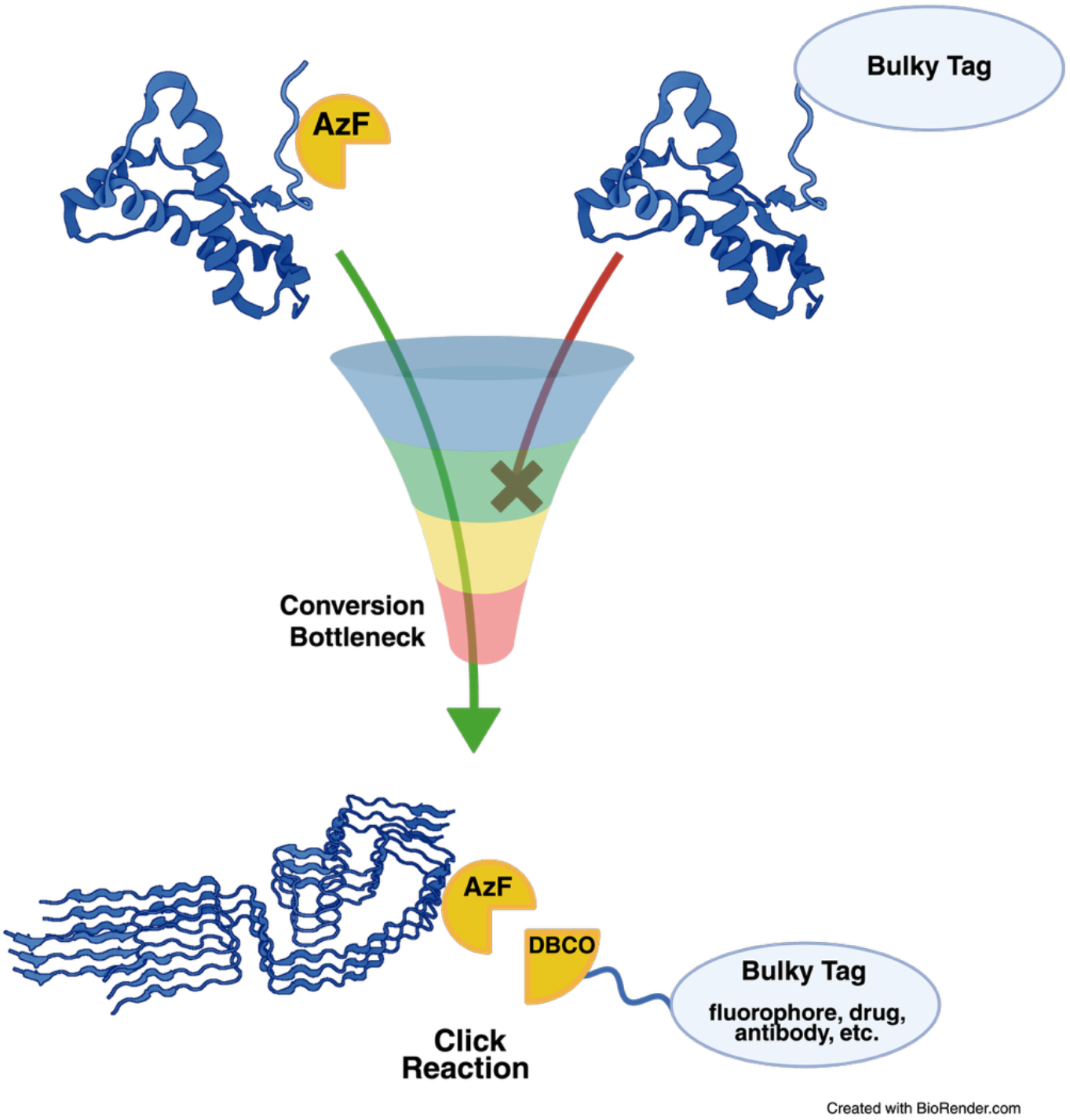

## Introduction

The prion protein (PrP) is a glycophosphatidylinositol (GPI)-anchored membrane glycoprotein that is enriched in neurons. It has the unorthodox ability to change from a properly folded cellular conformation termed PrP^C^ to various misfolded, protease-resistant, and infectious conformers collectively termed PrP^Sc1^. The presence of PrP^Sc^ can autocatalytically induce further misfolding of PrP^C^ in the brain, leading to a buildup of PrP^Sc^ resulting in fatal diseases such as Creutzfeldt-Jakob disease (CJD), bovine spongiform encephalopathy (BSE), and scrapie^2^. The initial misfolding of PrP^C^ can be triggered by exposure to infectious PrP^Sc^, genetic mutations in the PrP gene (*PRNP*), or spontaneous conversion of PrP^C3^. Cryo-electron microscopy (cryo-EM) studies reveal that PrP^Sc^ assembles into parallel in-register β-sheet (PIRBS) amyloid fibrils^4^. Many other functional and pathological amyloid proteins with prion-like self-propagation ability exist in species as diverse as yeast and humans^5, 6^. Notably, the propagation of α-synuclein and tau amyloid fibrils may play important roles in the pathogenesis of Parkinson’s Disease and Alzheimer’s Disease^78^.

To facilitate investigations into the structure, function, and cellular fate of PrP^Sc^ and other amyloid proteins, it would be useful to attach various reporters and ligands to specific sites of the amyloid protein of interest. However, amyloid formation is easily disrupted by perturbations in the precursor protein^9, 10^, making it challenging to introduce site-specific modifications into precursor proteins while maintaining the ability to form amyloid fibrils. We refer to this sensitivity as the conversion bottleneck.

In the absence of a general method to specifically modify amyloid proteins, various bespoke and traditional labeling techniques have been used. In the case of α-synuclein, a non-perturbing tetracysteine motif has been inserted into the protein to study fibril formation^11^. While this method works well for α-synuclein, it is not optimal for proteins that contain endogenous cysteines and disulfide bonds, such as PrP. Likewise, various sequences including His-tags, Myc-tags, and GFP-tags^10, 12, 13^ have been used to tag PrP. However, these large tags can only be accommodated in select locations and typically prevent the ability of PrP^C^ to convert into infectious PrP^Sc13^. An alternative approach has been to covalently attach fluorophores to WT PrP^Sc^ using chemically reactive groups such as NHS esters^14^. However, this approach is not site-specific since reactive compounds such as NHS esters covalently attach to multiple residues, thereby causing non-specific impacts on PrP^Sc^ structure and infectivity. These studies and others highlight the difficulties in specifically labeling PrP^Sc^ and other amyloid proteins specifically without disrupting their structure and function.

Many questions in prion biology remain unanswered due in part to our inability to produce specifically modified PrP^Sc^ molecules. For example, it is difficult to identify the specific structural motifs within the PrP^Sc^ replication interface that regulate infectious amyloid replication. Additionally, the PrP^C^ interactome has been determined^15, 16^, but the cellular interactome of PrP^Sc^ has not been determined. Finally, while several studies have identified critical elements of the prion neuroinvasion pathway^17^, our inability to produce labeled PrP^Sc^ inoculum makes it challenging to longitudinally track the route infectious prions take to invade the central nervous system from peripheral tissues *in vivo*. To answer these and other questions, it would be valuable to be able to specifically introduce a single conservative, chemically reactive group into PrP^C^ that doesn’t perturb the conformationally sensitive conversion process, which could then be subsequently modified in the PrP^Sc^ state.

Genetic code expansion has allowed for the selective substitution of non-canonical amino acids (ncAAs) into the primary amino acid sequence of a protein^18^. When labeling proteins, ncAAs can either be inserted as an already-functionalized reporter, such as a fluorescent ncAA, or as a small, non-functionalized reactive handle that allows for post-translational labeling. Substitution of fluorescent ncAAs has been previously used successfully to study α-synuclein, but in some cases altered aggregation kinetics^19^. We thought that using a smaller, more versatile handle would be more appropriate for labeling PrP^Sc^, so that the modified substrate could pass through the PrP^C^-to-PrP^Sc^ conversion bottleneck and then be used to specifically modify PrP^Sc^ post-conversion. We chose the click chemistry-amenable ncAA p-azido-L-phenylalanine (AzF), which can react bio-orthogonally with a chemically diverse library of commercially available alkyne- or dibenzocyclooctyne (DBCO)-containing reporters.

Our lab has developed an *in vitro* prion formation system in which recombinant PrP^C^ substrate is efficiently converted into PrP^Sc^ with high specific infectivity^20 21^. This system lends itself to metabolic incorporation of non-canonical amino acids (like AzF) into PrP^C^ during bacterial expression, followed by conversion of the ncAA-containing substrate into PrP^Sc^. Additionally, we can modulate the infectivity of the prions produced *in vitro* by either including or excluding phosphatidylethanolamine cofactor, producing infectious cofactor PrP^Sc^ or the non-infectious protein-only PrP^Sc^ conformers, respectively^20, 21^. This *in vitro* prion conversion system is both an ideal tool to produce selectively modified AzF prions and a tractable model to study the biochemical basis of prion infectivity. Therefore, we set out to test whether PrP^C^ containing a specific AzF substitution could pass through the conversion bottleneck in our *in vitro* conversion system to produce infectious prions amenable to click chemistry.

## Results and Discussion

### AzF-Substituted PrP^C^ can Convert into Self-Propagating PrP^Sc^

To make a clickable PrP substrate that could potentially be converted into PrP^Sc^, we made the conservative substitution of AzF for PrP residue W99 using amber codon reassignment in recoded B-95.ΔAΔ*fabR E. coli*^22^ and purified the resultant recombinant W99AzF PrP as described previously^23, 24^ (herein referred to as AzF PrP). To assess how the incorporation of AzF at position W99 of PrP^C^ affects *in vitro* prion conversion, we tested the ability of AzF PrP^C^ substrate to serially propagate cofactor PrP^Sc^ or protein-only PrP^Sc^ in continuous shaking reactions^25^. As determined by western blot, AzF PrP^C^ substrate successfully propagated both conformers without loss of conversion efficiency (**Figure 1 upper panel, lanes 9-11 and 12-14**). Furthermore, the proteinase K (PK)-resistant core of each conformer maintained its characteristic molecular weight over three rounds of propagation, demonstrating that AzF PrP^Sc^ faithfully templates prion conversion (**Figure 1 upper panel, compare lanes 9-11 with lanes 2-4 for protein-only PrP**^**Sc**^, **compare 12-14 with lanes 5-7 for cofactor PrP**^**Sc**^). Collectively, these results show that AzF PrP^C^ substrate can efficiently and faithfully propagate two different PrP^Sc^ conformations.

**Figure 1:**
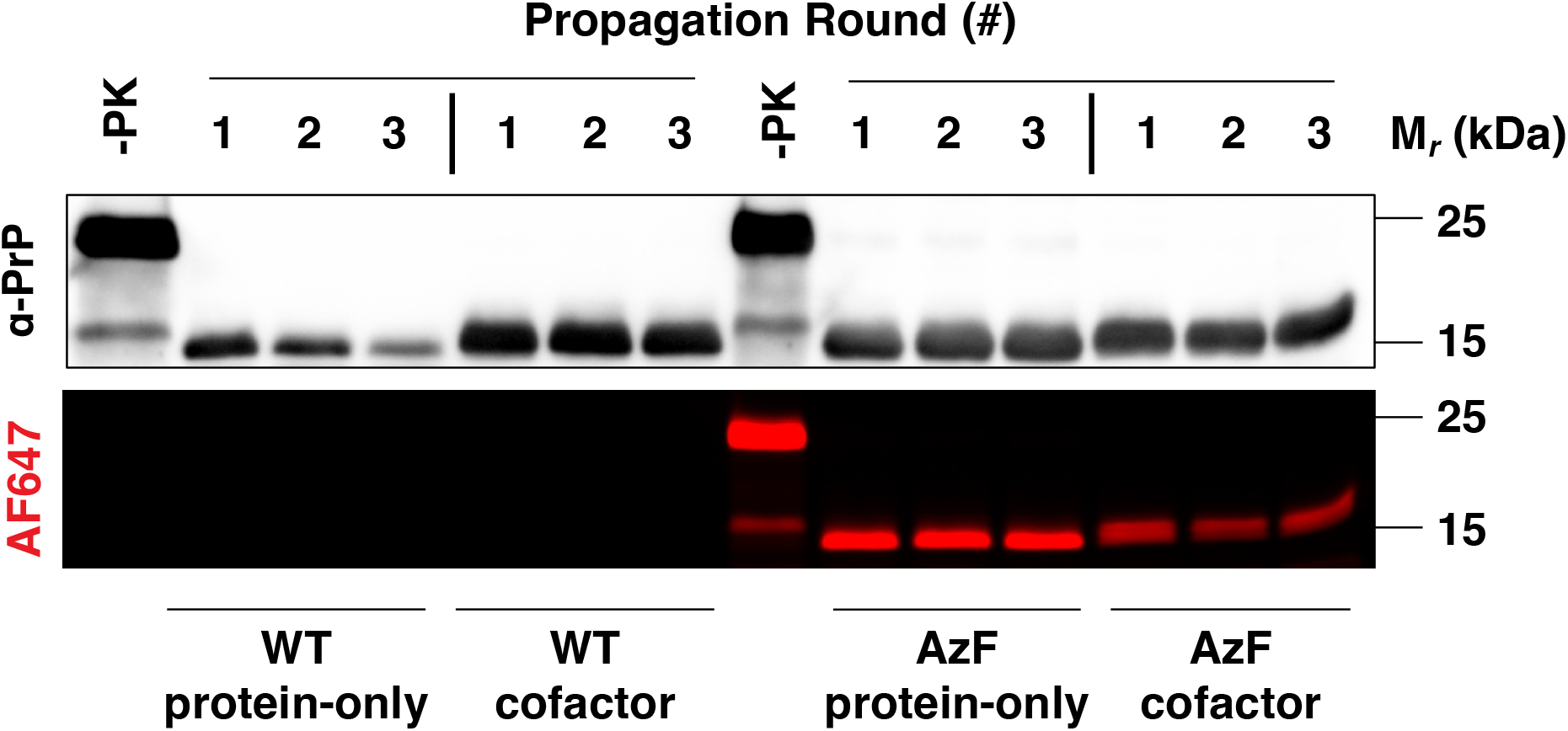
Effect of AzF Incorporation on Prion Strain Stability *in vitro*. Western blot (upper panel) and fluorescent gel (lower panel) showing three serial propagation rounds of WT protein-only PrP^Sc^, WT cofactor PrP^Sc^, AzF protein-only PrP^Sc^, and AzF cofactor PrP^Sc^, as indicated. All reactions were treated with PK except where otherwise indicated (-PK).

### AzF-Substituted PrP^Sc^ can be Clicked after Conversion

We next tested whether we could covalently attach a click reagent to newly formed AzF PrP^Sc^ molecules. To do this we selected incubated all the samples in Figure 1 with the large click ligand fluorophore AlexaFluor647-DBCO (AF647-DBCO) and used fluorescent gel imaging to detect AF647-labeled PrP. The results revealed that labeling was specific for AzF-containing PrP^Sc^, with no labeling observed for wild-type (WT) PrP^Sc^ (**Figure 1 lower panel, compare lanes 1-7 *versus* lanes 8-14**). Furthermore, clicking AF647-DBCO to AzF PrP^Sc^ does not change its PK-resistance (**Figure 1, lower panel lanes 8-14**). We compared the click efficiency of AF647-DBCO to AzF cofactor PrP^Sc^ and AzF protein-only PrP^Sc^ to the click efficiency of AzF PrP^C^ (which we used as an 100% reaction standard control). This analysis showed that the labeling efficiency of AF647-DBCO to AzF cofactor PrP^Sc^ was ∼50% and that of AzF protein-only PrP^Sc^ was ∼100% (**Supplemental Figure 1b**). These data show that both AzF PrP^Sc^ conformers in in their amyloid forms can be quickly and specifically reacted with a large click ligand.

### AF647-AzF PrP^Sc^ Can Seed PrP^Sc^ Formation

Given that we can covalently attach the bulky AF647-DBCO group to AzF PrP^Sc^, we next tested whether AF647-AzF PrP^Sc^ could act as a seed. To do this, we performed single round PrP^Sc^ conversion reactions using either AF647-AzF protein-only PrP^Sc^ or control WT protein-only PrP^Sc^ as seeds and either AzF PrP^C^ or WT PrP^C^ as substrates. Western blot analysis of the reaction products shows the presence of PK-resistant PrP^Sc^ molecules in reactions seeded with AF647-AzF PrP^Sc^ seed **(Figure 2, top panel, lanes 2 and 4)**. To confirm the formation of new PrP^Sc^ molecules from AzF PrP^C^ substrate, we treated reaction products with BODIPY-DBCO and then used fluorescent gel imaging to discriminate between AF647-AzF PrP^Sc^ seed (**Figure 2, middle panel, lanes 1-4**) and BODIPY-AzF PrP^Sc^ product. The results confirmed that AF647-AzF PrP^Sc^ can efficiently seed conversion of AzF PrP^C^ substrate into new AzF PrP^Sc^ molecules, as evidenced by a robust PK-resistant PrP^Sc^ band with a strong BODIPY signal **(Figure 2, bottom panel, lane 2)**. Moreover, AF647-AzF PrP^Sc^ appeared to seed the formation of new PrP^Sc^ molecules just as efficiently as WT PrP^Sc^ molecules **(Figure 2, bottom panel, compare lanes 2 versus lane 6)**. Control reactions using WT PrP^C^ substrate showed no BODIPY fluorescence **(Figure 2, bottom panel, lanes 3-4 and 7-8)**. Taken together, these results show that AF647-AzF protein-only PrP^Sc^ maintains the ability to seed PrP^Sc^ conversion reactions, despite the presence of a bulky fluorophore at position 99.

**Figure 2:**
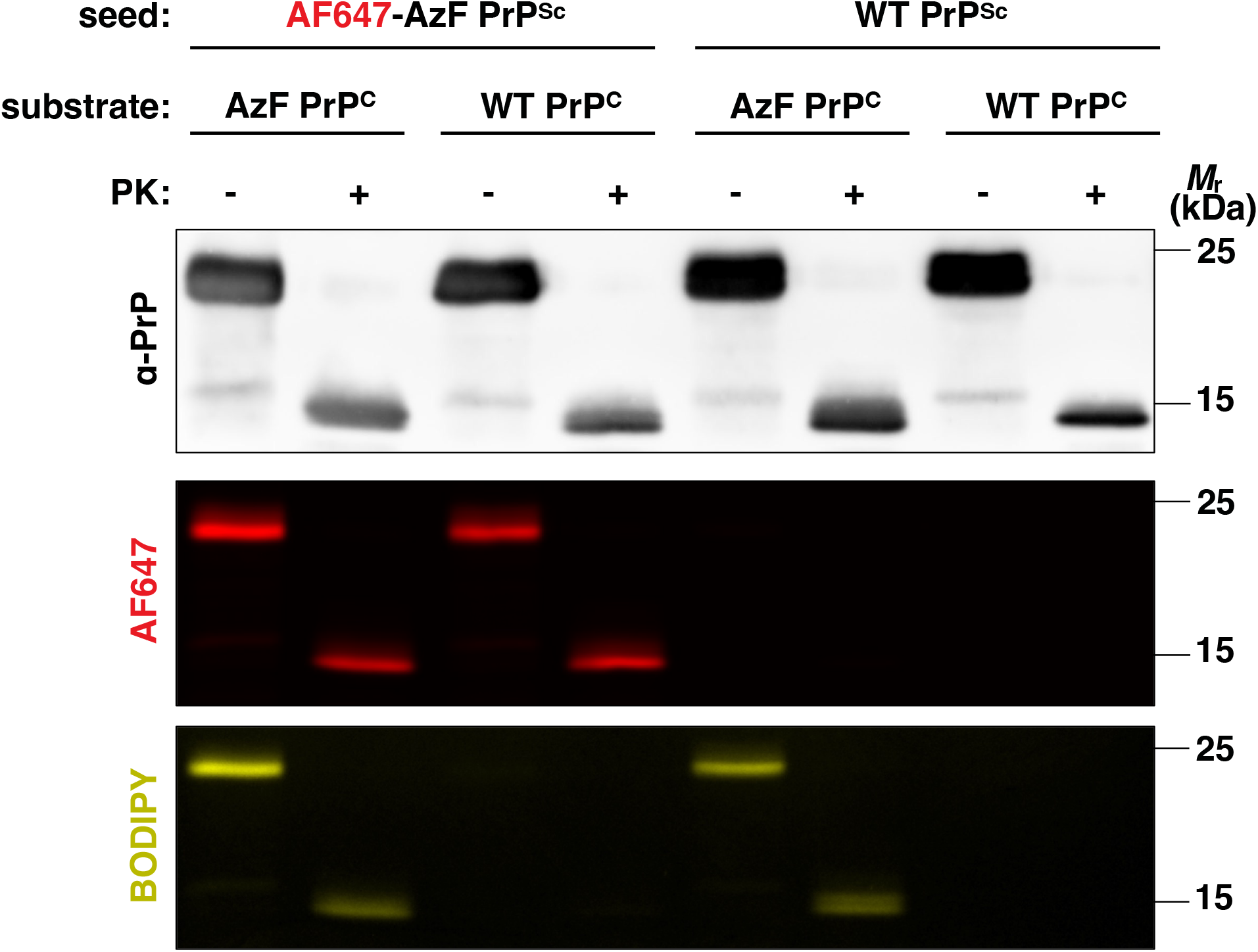
Seeding Ability of AF647-AzF PrP^Sc^. Western blot (top panel) and fluorescent imaging (middle panel = AF647; bottom panel = BODIPY) of single-round conversion reactions. AzF PrP^C^ or WT PrP^C^ substrates were seeded with either AF647-AzF protein-only PrP^Sc^ or WT protein-only PrP^Sc^ as indicated, and all products were subsequently labeled with BODIPY-DBCO. Samples were treated with PK where indicated.

### AzF cofactor PrP^Sc^ and AF647-AzF cofactor PrP^Sc^ are Infectious

To assess the infectivity of AzF PrP^Sc^ and AF647-AzF PrP^Sc^ molecules, we serially propagated AzF cofactor PrP^Sc^, AF647-AzF cofactor PrP^Sc^, and AzF protein-only PrP^Sc^ for 26 rounds to dilute out the original round 1 seeds beyond Avagadro’s limit. We then inoculated M109 bank vole PrP knock-in (kiBVM) mice^16^ with the products from serial propagation round 26. Mice inoculated with either AzF cofactor PrP^Sc^ or AF647-AzF cofactor PrP^Sc^ developed clinical signs scrapie with 100% attack rate and incubation times of 197 days (s.d.=5, n=8) and 185 days (s.d.=14, n=8), respectively (**Figure 3, yellow squares and red inverted triangles**), similar to positive control mice inoculated with WT PrP^Sc^ **(Figure 3 blue triangles)**. In contrast, mice inoculated with AzF protein-only PrP^Sc^ did not develop scrapie, as shown by the ongoing (>450 days) survival of these mice (**Figure 3, brown squares**), similar to negative control mice inoculated with unconverted reaction cocktail **(Figure 3, black crosses)**.

**Figure 3:**
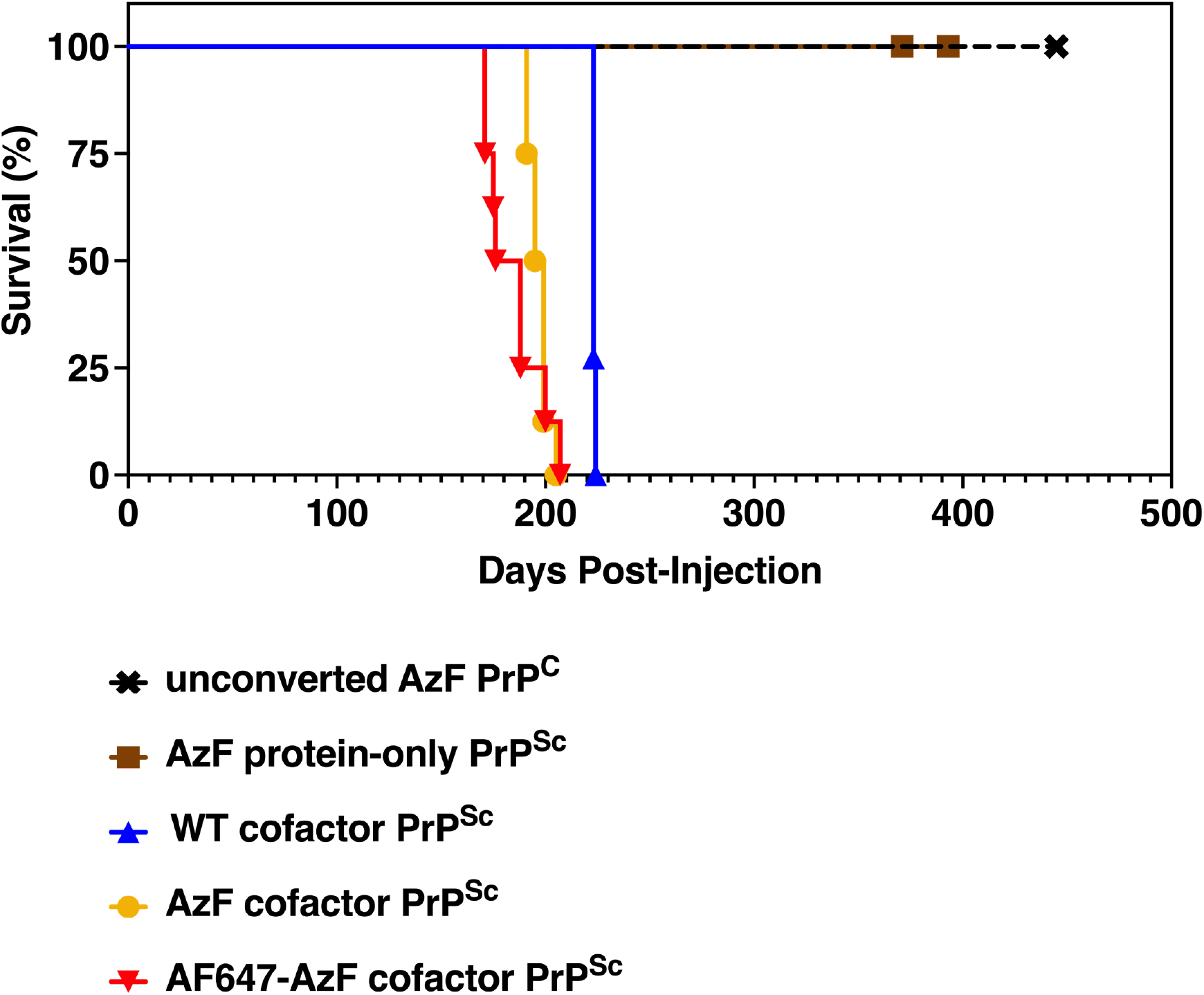
Bioassay of AzF PrPSc and AF647-AzF PrP^Sc^ Infectivity. Survival curves of kiBVM mice inoculated with various serially propagated PrP^Sc^ products, as indicated (n=7-8 in each group). Mice were monitored daily and sacrificed upon developing clinical signs of scrapie.

Post-mortem, we also examined neuropathology and biochemical properties of the resultant brain-derived PrP^Sc^. Microscopic examination revealed extensive vacuolation in the brains of mice inoculated with AzF cofactor PrP^Sc^ and AF647-AzF cofactor PrP^Sc^, consistent with scrapie infection (**Figure 4, see bottom two panels**). Furthermore, the degree of vacuolation was similar between these two conformers and WT cofactor PrP^Sc^ across various brain regions examined (**Supplemental**, **compare red circle and yellow triangles with blue diamonds**). Taken together, these data show that AzF cofactor PrP^Sc^ and AF647-AzF cofactor PrP^Sc^ are infectious and display similar patterns of neurotropism.

**Figure 4:**
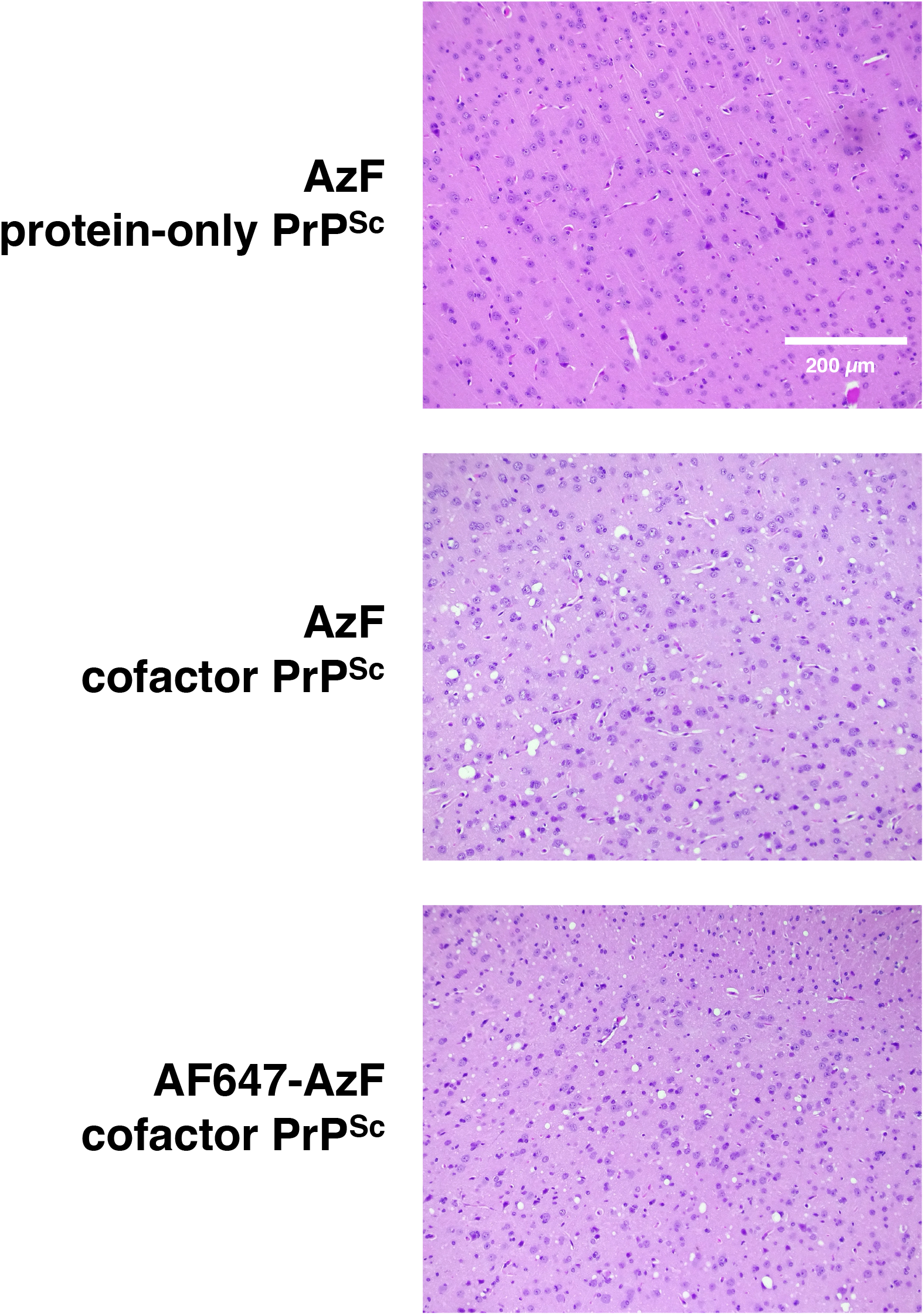
Neuropathology of Inoculated Mice. Representative hematoxylin and eosin (H&E) stained microscopic images of the cerebral cortex in mice inoculated with AzF protein-only PrP^Sc^, AzF cofactor PrP^Sc^, or AF647-AzF cofactor PrP^Sc^, as indicated.

To directly measure the accumulation of PrP^Sc^ in the brains of mice that succumbed to scrapie, we performed western blots of brain homogenates. The results showed extensive PK-resistant PrP^Sc^ formation in the brains of mice inoculated with AzF cofactor PrP^Sc^ and AF647-AzF cofactor PrP^Sc^ **(Figure 5, lanes 8 and 10)**, with glycoform profiles similar to that of mice inoculated with WT cofactor PrP^Sc^ (**Figure 5, compare lanes 8 and 10 to lane 12**). In contrast, no PrP^Sc^ was detected in mice inoculated with AzF protein-only PrP^Sc^, negative control mice inoculated with unconverted AzF PrP substrate, or age-matched uninoculated kiBVM mice **(Figure 5, lanes 2, 4, and 6)**. Taken together, these data suggest that AzF-substituted prions are infectious, self-propagate *in vivo*, and retain infectivity when conjugated to a model click ligand.

**Figure 5:**
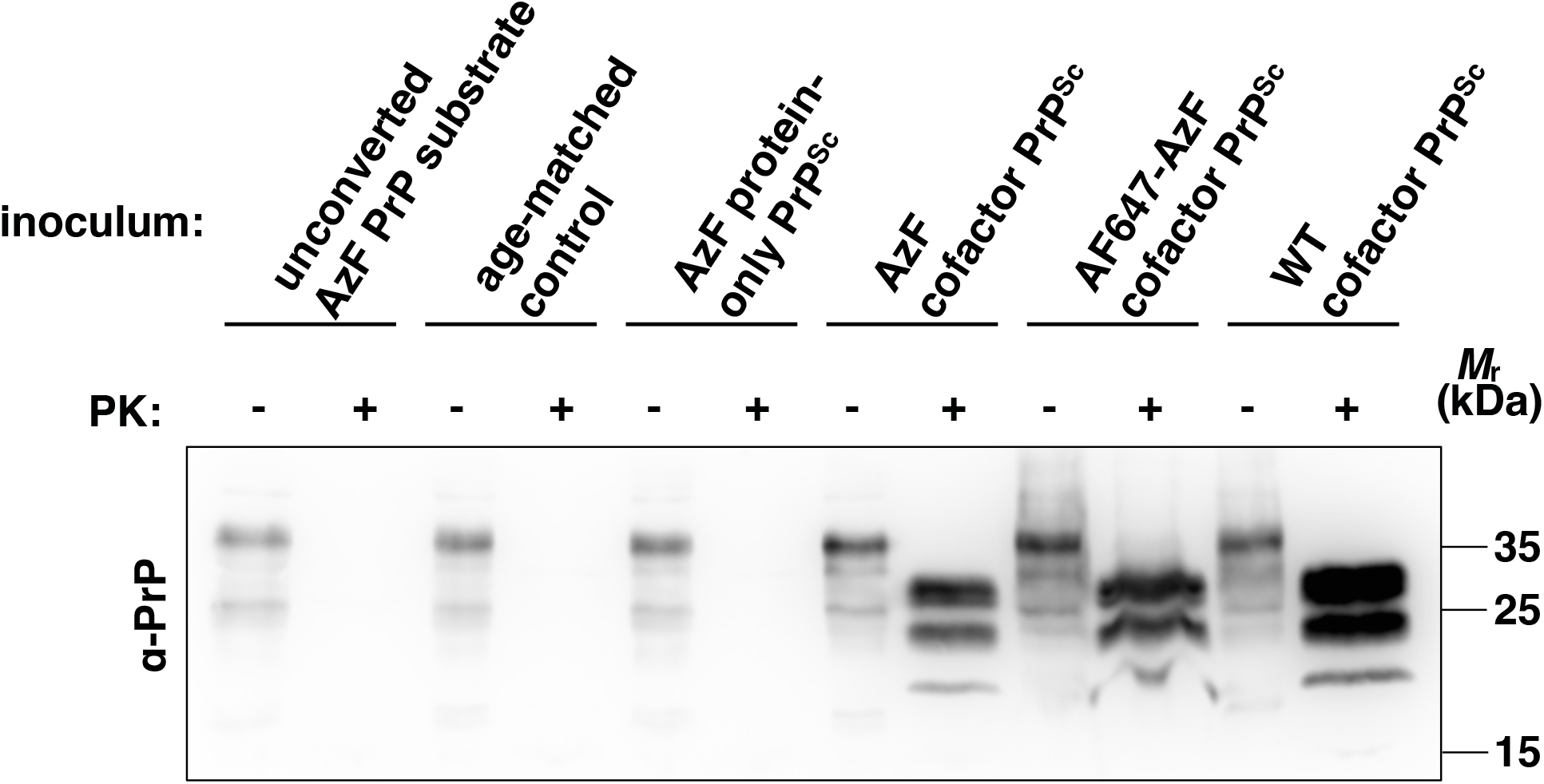
*In vivo* Accumulation of PrP^Sc^ in Brains of Inoculated Mice. Western blot of brain homogenates from kiBVM mice inoculated with various serially propagated PrPSc products, as indicated. Samples were treated with PK where indicated.

### Visualization of Near-Infrared-Labeled PrP^Sc^ in Intact Mouse Brain

In order to explore the possible uses of click-conjugated PrP^Sc^, we clicked the near-infrared (NIR) dye Cy7.5-DBCO to AzF protein-only PrP^Sc^ **(Figure 6a)** and tested its application as a tool for real-time *in situ* visualization of prions inoculated into animals. We injected Cy7.5-conjugated PrP^Sc^ intracerebrally into a mouse at two independent injection sites ∼5 mm apart and employed NIR imaging to determine whether the two inoculation sites could be visualized and resolved. NIR imaging of the intact mouse brain revealed two well-resolved puncta of fluorescent PrP^Sc^, corresponding to each site of Cy7.5-PrP^Sc^ injection (**Figure 6b, see arrows**). These data show that NIR conjugated PrP^Sc^ can be used to visualize and track inoculum in intact tissue.

**Figure 6:**
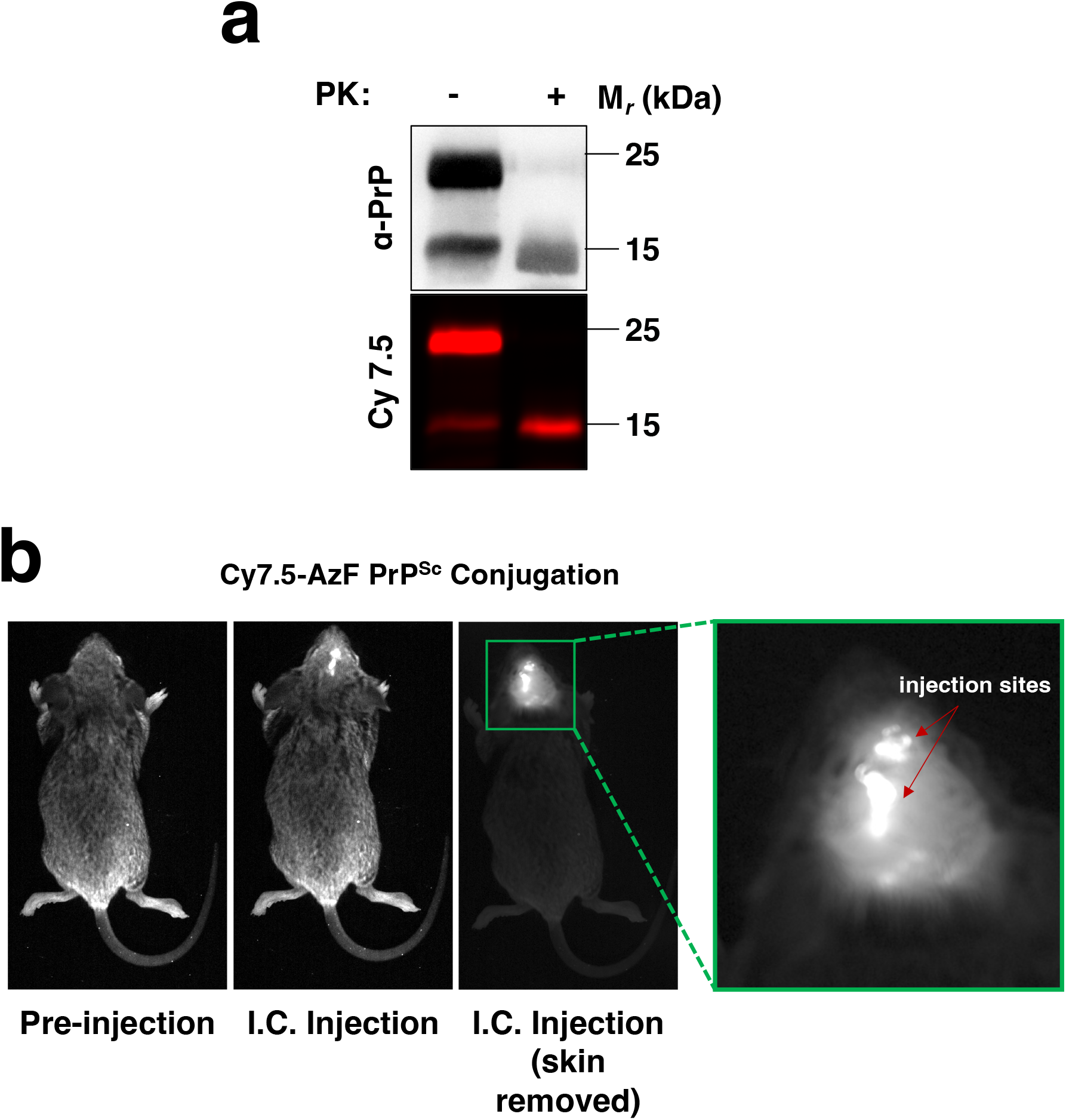
Imaging NIR-Labeled PrP^Sc^ in an Intact Mouse Brain. (A) Fluorescent gel and western blot of AzF PrP^Sc^ clicked with the NIR dye Cy7.5-DBCO. Samples were treated with PK where indicated. (B) Fluorescent image of mouse injected twice intracerebrally with Cy7.5-PrP^Sc^ resuspended in 1X PBS (15 µL total, 8.9 μM). Blue stars indicate independent injection sites.

### Potential Applications

AzF cofactor PrP^Sc^ can be immediately used to study various properties of infectious prions by clicking to various ligands. For instance, NIR-labeled PrP^Sc^ molecules can be used to glean information about how peripherally administered scrapie invades the central nervous system and where scrapie localizes during initial infection. Additionally, by monitoring for NIR signal disappearance from an animal, AzF prions can be used to measure the lifespan of and initial inoculum *in vivo*. In trying to understand prion trafficking *in vivo*, previous studies have had to sacrifice a different animal for each timepoint during a prion infection^26^. Significantly, NIR-labeled PrP^Sc^ could be used to track the fate of prions injected into a single animal *in situ* longitudinally.

AzF cofactor PrP^Sc^ could also be used to study the behavior of prions in cells. For instance, the AzF handle could be used to covalently immobilize AzF prions crosslinked to potential interactors from cell lysates to determine the cellular interactome of exogenously applied PrP^Sc^.

The AzF handle also provides a unique tool for the precise manipulation of prions to study the relationship between prion structure and properties such as seeding and infectivity. Utilizing a library of prions with AzF moved throughout the structure, the contribution of each position to key biochemical properties can be isolated.

The breadth of click ligands presents the opportunity for more applications than the few described here. Some additional ligands include antibodies, membrane-targeting lipids, E3 ligases, small molecule drugs, and FRET-compatible fluorophores.

## Limitations

Although the generation of infectious prions amenable to click chemistry represents an important technical advance, there are several limitations of our study. One limitation is that we only tested one residue for AzF substitution. Based on the general similarity of their side chain structures, we expect that substitution of many other aromatic residues within the PrP sequence by AzF would also overcome the conversion bottleneck, but it is possible that AzF substitution of some residues would not be tolerated. In any case, W99AzF cofactor PrP^Sc^ itself can be clicked to many different ligands and used for multiple types of studies, such as tracking inoculum in an animal, selective pulldown of infectious AzF inoculum, and fibril replication studies.

Another limitation is that our *in vitro* propagation system with recombinant PrP substrates cannot, at this time, be used for propagating natural prion strains^21, 25^. Replication of natural prion strains requires propagation with native PrP^C^ molecules in animals, cells, or brain homogenate Protein Misfolding Cyclic Amplification (PMCA) reactions in in order to retain strain properties^21, 27^. However, site-specific ncAA incorporation in mammalian cells is possible^28^, providing an opportunity to make native AzF prions with natural strain properties. Although our system is only suitable for recombinant prions, it produces PrP^Sc^ with high conversion efficiency and with a high level of specific infectivity comparable to natural prion strains^21, 29^.

Finally, we observed ∼50% reaction of W99AzF cofactor PrP^Sc^ with AF647-DBCO. This incomplete reaction limits our ability to measure the infectivity of AF647-AzF cofactor PrP^Sc^ independent of unreacted W99AzF cofactor PrP^Sc^. Due to this limitation, we did not attempt to determine the specific infectivity of AF647-AzF cofactor PrP^Sc^ by end-point titration^30^. Given that click chemistry typically results in quantitative labeling and that we obtained ∼100% reaction of W99AzF protein-only PrP^Sc^ with AF647-DBCO, we speculate that incomplete labeling of AzF cofactor PrP^Sc^ fibrils is not likely due to a technical issue, but rather a consequence of the specific conformation around residue 99 of W99AzF cofactor PrP^Sc^. The repeating nature of prion fibrils could cause neighboring AF647-DBCO groups to sterically and electrostatically clash, which provides a mechanism for varying levels of reaction efficiency between protein-only and AzF cofactor PrP^Sc^. In the case of W99AzF cofactor PrP^Sc^, it is likely that only one bulky AF647-DBCO group can be accommodated in the space between two rungs of the amyloid ladder. Indeed, click reaction efficiency could potentially be used to probe the intermolecular structure of PrP^Sc^ fibrils. While complete click reaction is necessary for certain experimental designs, it is not required for experiments such as tracking, selective pulldown, and imaging.

## Conclusion

In this paper we report the first successful production of click chemistry-reactive infectious prions. Substitution of PrP residue W99 with AzF effectively smuggles click potential through the conformationally-sensitive conversion process, which has historically stood as a roadblock to converting various modified PrP^C^ substrates into PrP^Sc^. In bypassing the conversion bottleneck, large or otherwise disruptive groups can subsequently be directly attached by click chemistry to infectious AzF PrP^Sc^, making regions that previously were not susceptible to modification available for study.

In addition to being used for studying PrP^Sc^, AzF incorporation could be used to answer longstanding questions about other functional or disease-relevant amyloids^5^. For example, both tau and α-synuclein can be converted from recombinant precursor proteins into a variety of different conformers *in vitro* ^31^. AzF incorporation is an attractive strategy for investigating how specific structural determinants of tau and α-synuclein amyloid fibrils contribute to strain properties. More generally, our results suggest that AzF incorporation provides a uniquely powerful approach to studying the biology of infectious prions and other amyloid proteins.

## Materials and Methods

### Ethics Statement

All animals were housed and cared for according to standards set by the Guide for the Care and Use of Laboratory Animals of the National Research Council. All mouse experiments were conducted in accordance with protocol supa.su.1, as reviewed and approved by Dartmouth College’s Institutional Animal Care and Use Committee, operating under the regulations/guidelines of the NIH Office of Laboratory Animal Welfare (assurance number A3259-01).

### Cloning of AzF PrP

Recombinant bank vole M109 W99AzF PrP, hereafter referred to as AzF PrP, was generated through Gibson assembly. An expression construct was made using pET-22b(+) (Sigma-Aldrich, St. Louis, MO) cut with NdeI/HIndIII (New England Biolabs, Ipswich, MA), gel-purified, and used for Gibson Assembly (New England Biolabs, Ipswich, MA) with the bank vole amber-codon substituted (bolded in sequence below) PrP oligo (IDT, Newark, NJ), codon-optimized for expression in bacteria:

5’-TTGTTTAACTTTAAGAAGGAGATATACATATGAAAAAGCGTCCAAAGCCAGGTGGCTGGAATACGGGGGGTAGTCGTTATCCTGGTCAGGGCTCCCCGGGCGGTAATCGCTATCCGCCGCAAGGCGGGGGTACATGGGGCCAACCACACGGTGGCGGTTGGGGCCAACCTCACGGGGGCGGTTGGGGGCAACCTCATGGCGGTGGGTGGGGTCAACCGCATGGGGGTGGCTGGGGT CAAGGTGGCGGGACTCACAATCAG**<u>TAG</u>**AACAAGCCTTCTAAACCGAAAACGAATATGAAG CACGTTGCGGGTGCAGCCGCTGCGGGTGCTGTTGTAGGTGGCTTAGGTGGCTACATGCTGGGTTCTGCCATGAGTCGCCCGATGATTCACTTCGGTAATGATTGGGAAGATCGTTATTATCGCGAGAATATGAATCGTTACCCAAATCAGGTCTACTATCGTCCGGTAGACCAGTACAACAATCAGAATAATTTTGTACATGATTGCGTTAATATCACGATTAAACAACATACGGTTACTACTACCACGAAGGGCGAGAATTTTACCGAGACCGACGTCAAGATGATGGAACGCGTAGTCGAGCAAATGTGTGTAACACAGTATCAAAAGGAGAGTCAAGCCTACTATGAGGGTCGCTCATCCTAATAAAAGCTTGCGGCCGCACTCGAGCACCACCACC-3’

The resultant plasmid was transformed into DH5a cells (Invitrogen, Waltham, MA), mini-prepped (Qiagen, Hilden, Germany), and confirmed by sequencing.

### Expression of Recombinant AzF PrP

To express AzF PrP, B-95.ΔAΔ*fabR E. coli*, provided by the RIKEN BRC through the National BioResource Project of the MEXT, Japan (cat. RDB13712), were co-transformed with the AzF PrP plasmid and the pEvol-pAzFRS.2.t1 expression plasmid (Addgene, Watertown, Ma), which codes for the tRNA synthetase/tRNA pair necessary for p-azido-L-phenylalanine (AzF) incorporation^22^. Double transformants were selected on LB agar plates and grown overnight in 2x Yeast Tryptone (2x YT) (Sigma-Aldrich, St. Louis, MO) media at 37°C. Overnight culture was centrifuged (4200 x *g*, 10 min) and resuspended in 3mL of fresh 2x YT media without antibiotics and used to inoculate 500mL of 2x YT media. Once the OD_600nm_ reached 1-3, arabinose was added to a final concentration of 0.2% w/v, AzF was added to a final concentration of 1 mM, and bacteria were incubated for 1 hr at 37°C to induce expression of the tRNA/pAzFRS synthetase pair. After 1 hr, protein expression was induced by adding IPTG to a final concentration of 1 mM and bacteria were grown for 8-16 hrs at 30°C. Bacteria were pelleted at 16,000 x *g* for 10 min.

All steps were done in the presence of 34 µg/mL chloramphenicol and 50 µg/mL ampicillin, unless otherwise noted. All incubation steps were carried out at 230 rpm in a VWR 1585R shaking incubator (VWR, Radnor, PA).

### Purification of AzF PrP

AzF PrP was purified using the PrP purification protocol previously described by Makarava and Baskakov, 2008^23^. In brief, bacteria expressing W99AzF PrP were harvested by centrifugation. Cell pellets were resuspended in Bugbuster (MilliporeSigma, Burlington, MA) containing Lysonase (MilliporeSigma, Burlington, MA) and EDTA-free protease inhibitor (Roche, Indianapolis, IN) and intermittently sonicated for 30 sec every 5 min for 30 min total using a handheld sonicator at setting 8.5 (Sonopuls Amtrex, St-Laurent, QC). Inclusion bodies containing PrP were isolated by centrifuging cell lysate at 16,000 x *g* for 20 min and resuspending with BugBuster a total of three times. PrP inclusion bodies were solubilized in a buffer containing urea and PrP was purified through nickel-affinity chromatography, followed by size-exclusion chromatography. Reverse-phase (C4) HPLC (Sepax Technologies, Inc., Newark, DE) was performed to yield pure AzF PrP. PrP eluate from the C4 column was lyophilized and frozen at -20°C for long-term storage. Importantly, no glutathione was used in the purification process due to its capacity to reduce the azide group.

### *In vitro* Conversion of AzF PrP^C^ into PrP^Sc^

Prior to use, lyophilized AzF PrP^C^ was resuspended in water to a concentration of 0.12 mg/mL and was used to make substrate cocktail for PrP^Sc^ conversion as described by Walsh *et al* 2023^25^. In brief, the substrate cocktails contain PrP (6ug/mL) and reaction buffer composed of: NaCl (135mM), EDTA (5mM), Triton X-100 (0.15%), Tris buffer (20mM, pH 7.5) and lipid cofactor (optional, 10-20%). *In vitro* conversion reactions were initiated by adding 100uL of 6 µg/mL PrP^Sc^ (in reaction buffer) to 400µL of substrate shaking cocktail and shaken for 72 hr. For serial propagations, 100uL of the newly converted PrP^Sc^ was used to seed a new 400uL shaking cocktail to start the next round of serial propagation. AzF cofactor PrP^Sc^ was produced through seeding cofactor-containing AzF PrP^C^ substrate cocktail with WT cofactor recPrP^Sc^. AzF protein-only PrP^Sc^ was produced by seeding AzF PrP^C^ substrate cocktail with WT cofactor recPrP^Sc^ and withdrawing cofactor over ∼30 rounds of propagation as previously described^21^.

All propagation reactions were performed while continuously shaking for 72 hr at 37°C at 2,000 rpm in a 3 mm orbit Ohaus shaker (Parsippany, NJ).

### Click Reactions with AzF PrP

After PrP^Sc^ propagation, the *in vitro* prion conversion reaction was centrifuged and the supernatant was aspirated to remove soluble PrP^C^. The PrP^Sc^ pellet was sonicated for 30 sec, 160W, at room temperature in a QSonica sonicator to break up the pellet (Misonix Model 3000, Farmingdale, New York). DBCO-dyes were resuspended according to manufacturer recommendations (AF647-DBCO, JenaBioscience, Thuringia, Germany; BODIPY-FL-PEG4-DBCO, JenaBioscience, Thuringia, Germany; Cy7.5-DBCO, RuixiBio, Xi’ An City, China). Click reactions were performed with an AzF PrP^Sc^ concentration of 30 µg/mL and a 20-fold molar excess of DBCO-dye in a total volume of 100 µL. The click reaction was allowed to proceed for 1 hr at 25°C while shaking at 750 rpm (Ohaus, Parsippany, NJ). To remove excess DBCO-dye, the reaction was diluted 10-fold with 1X PBS + 0.1% Triton X-100, sonicated, centrifuged, and aspirated. This washing process was repeated a total of three times to yield a washed protein pellet.

All centrifugation steps were done at 18,000 x *g* for 30 min at 25°C.

## Acknowledgements

This study was funded by the National Institute for Neurological Diseases and Stroke (1R37NS125431, R01NS117276 and R01NS118796 to S.S.) and the National Institutes of Health (P20-GM113132 to Dean Madden). We would like to thank Dr. Daniel Walsh for helpful advice on protein purification protocols. We would also like to thank Dr. Kensaku Sakamoto from the RIKEN Center for Life Science Technologies for providing the B-95.ΔAΔ*fabR E. coli*.

