## Supplementary Figures and Methods for "Generation of Infectious Prions Amenable to Site-specific Click Chemistry"

**Proteinase K Digestion and PrP$^{Sc}$ Visualization:**

To detect the presence of protease-resistant PrP$^{Sc}$, samples were subjected to Proteinase K (PK) digestion and analyzed by western blotting, as described in Walsh *et al*. 2023[1]. Brain-derived samples underwent additional processing including bead mill homogenization and centrifugation at 200 x *g* for 30 seconds to remove brain debris and to isolate the PrP$^{Sc}$-containing supernatant for PK digestion. In brief, samples were incubated with 20 µg/mL of PK for 30 min at 37℃ at 750 rpm in a 3 mm orbit Ohaus shaker (Parsippany, NJ). After digestion, samples were quenched with 4 mM phenylmethylsulfonyl fluoride (PMSF), boiled in Laemmli SDS loading buffer (Bioland Scientific, Paramount, CA) with 2-Mercaptoethanol for 10 minutes at 95℃ and run on a 12% polyacrylamide gel. Protein was transferred to a polyvinylidene fluoride (PVDF) membrane and was blotted as described previously with mAb 27/33 (epitope: 142-149, mouse numbering) and horseradish peroxidase-linked sheep anti-mouse antibodies[2]. Fluorophore conjugated PrP$^{Sc}$ was imaged after SDS-PAGE using an Azure 600 bioimager (Azure Biosystems, Dublin, CA). The following excitation and emission settings were used to acquire images for each fluorophore: AF647, Ex. 628/16 nm, Em. 684/12 nm; BODIPY, Ex. 472/15 nm, Em. 513/8.5 nm; Cy 7.5, Ex. 784 nm, Em. 832/18.5 nm.

**Preparation of Inoculum:**

PrP$^{Sc}$ propagation reactions were washed of unconverted PrP$^{C}$ by centrifuging for 30 min at 18,000 x *g* and aspirating the supernatant. After washing, inoculum was diluted 1:10 in 1X PBS + 1% bovine serum albumin. *In vivo* inoculation was performed as described in Piro *et al.* 2009[2]. For Cy7.5-PrP$^{Sc}$ injection, protein was concentrated to 8.9 µM and a total volume of 15uL was injected intracerebrally, divided between two sites.

**Scrapie Inoculation and Diagnosis:**

Knock-in female mice expressing bank vole M109 PrP (termed kiBVM mice)[3] between 4-5 weeks old were intracerebrally injected with 30 µL inoculum at 0.6 µg/mL PrP. Scrapie symptoms were monitored daily and a clinical diagnosis of scrapie was made based on the onset of wide gait, shaking, and/or circling[4]. Data was analyzed using GraphPad Prism 10; time to onset was reported as mean and standard error of the mean (SEM).

**Neuropathology:**

Within 24 hrs of scrapie symptom onset, sick mice were sacrificed and whole mouse brains were harvested and sliced in half parasagittally. For Western blot samples, ½ brains were frozen at -80℃ until western blot sample preparation[2]. For pathology samples, ½ brains were placed into 10% formalin for fixation, decontaminated by immersion in 88% formic acid for 1 hr, paraffin embedded, sliced parasagittally, and stained with hematoxylin and eosin by the Dartmouth Hitchcock Research Pathology Service Core (Lebanon, NH). Tissue samples were scored between 0-5 as previously described[5].

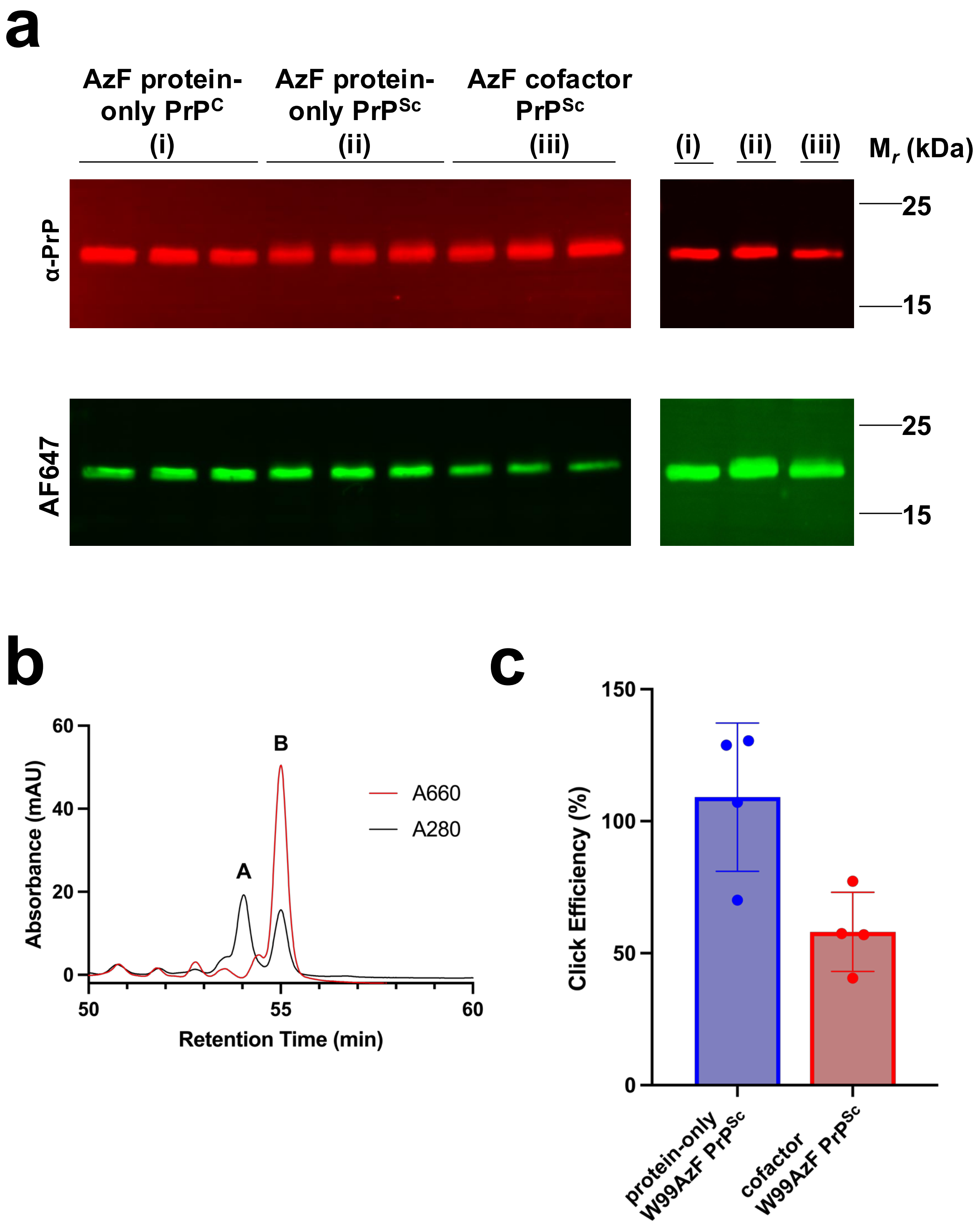


**Supplemental Figure 1: Determination of Click Efficiency of Protein-only and Cofactor AzF PrP$^{Sc}$.**

(A) 3 µg of either AzF $PrP^{C}$ (i), AzF protein-only $PrP^{Sc}$ (ii), or AzF cofactor $PrP^{Sc}$ (iii) was reacted with 20 µM AF647-DBCO for 1.5 hours, 25°C, 750 rpm (Ohaus, Parsippany, NJ). The reaction was quenched with a final concentration of 100 mM sodium azide. Samples were boiled in 1X Laemmli SDS loading buffer (Bioland Scientific, Paramount, CA) with 2-Mercaptoethanol for 10 minutes at 95°C and run on a 12% polyacrylamide gel. Protein was transferred to a polyvinylidene fluoride (PVDF) membrane and was blotted as described previously with mAb 27/33 (epitope: 142-149, mouse numbering) and Amersham CyDye800 goat anti-mouse (Cytiva, Marlborough, MA) antibodies to quantify total PrP (Ex. 784 nm, Em. 832/18.5 nm). AF647-PrP conjugation was directly measured using AF647 fluorescence (Ex. 628/16 nm, Em. 684/12 nm) on an Azure 600 bioimager (Azure Biosystems, Dublin, CA). All samples are technical replicates.

(B) Quantification of click reaction efficiency for AzF $PrP^{C}$ with AF647-DBCO using analytical HPLC. AF647-AzF $PrP^{C}$ (14.4 µg) was prepared as described in the materials and methods, and the reaction was quenched with excess sodium azide. This clicked $PrP^{C}$ was combined with unclicked AzF $PrP^{C}$ (21.6 µg), such that the final concentration was 40% clicked $PrP^{C}$, 60% unclicked $PrP^{C}$. The protein sample was denatured in 2M urea and loaded onto a C4 reverse phase column (Sepax Technologies, Inc., Newark, DE). Reverse phase chromatography was performed as described in Makarava and Baskakov, 2008[6] to resolve clicked $PrP^{C}$ from unclicked $PrP^{C}$. Both PrP species eluted at ~32% acetonitrile, typical for PrP[6]. Unclicked PrP (Peak A) was confirmed by 280nm absorbance and a clear lack of 660nm absorbance. AF647-AzF PrP (Peak B) was confirmed by its large 660nm absorbance, consistent with AF647 conjugation. After integrating peaks A and B, the total protein composition was 40.27% peak A, 59.73% peak B, which is as expected if AF647-DBCO clicked quantitatively to AzF $PrP^{C}$ in the initial reaction to create the clicked-PrP peak B. The congruence between input protein ratio and the HPLC peak integrations suggests a quantitative reaction between AzF $PrP^{C}$ and AF647-DBCO.

(C) The percent of total AzF PrP clicked to AF647-DBCO was determined by dividing the AF647 fluorescence (**A**, bottom panel) by total PrP signal (**A**, top panel). The AF647/CyDye800 ratio for AzF protein-only $PrP^{Sc}$ and AzF cofactor $PrP^{Sc}$ was normalized by setting the average AF647/CyDye800 ratio for AzF $PrP^{C}$ to 100%, consistent with the click reaction for AzF $PrP^{C}$ being quantitative. Normalized, technical quadruplicate data was plotted as mean and standard error of the mean using GraphPad Prism 10 (GraphPad, San Diego, CA).

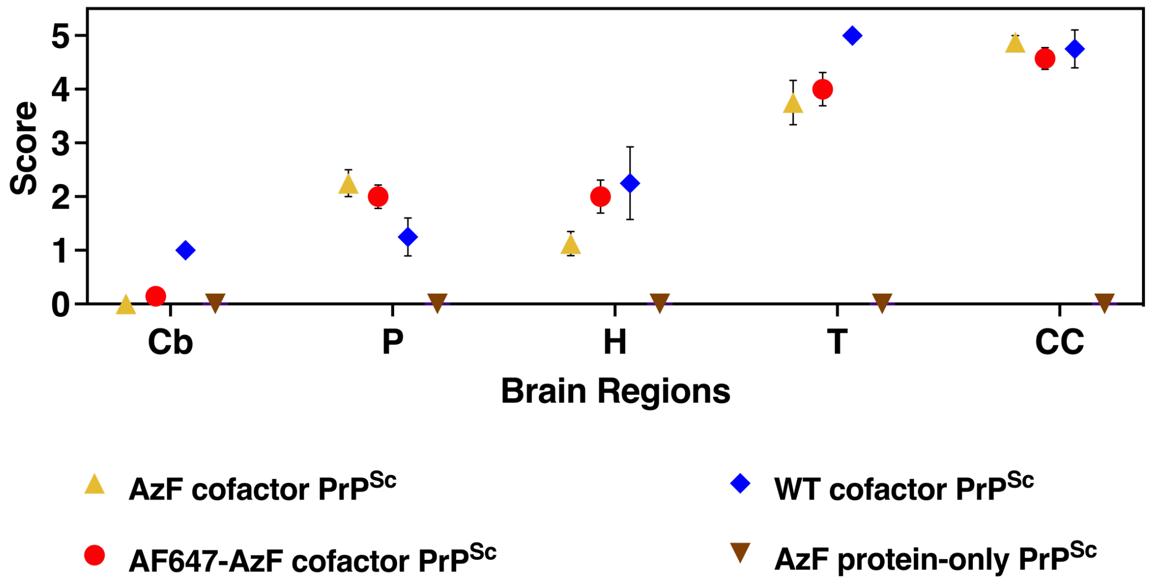


**Supplemental Figure 2: Neuropathology of Mice Inoculated with AzF $PrP^{Sc}$ and AF647-AzF $PrP^{Sc}$**
Representative hematoxylin and eosin (H&E) stained images showing vacuolation in the cerebral cortex of mice inoculated with AzF protein-only $PrP^{Sc}$, AzF cofactor $PrP^{Sc}$, or AF647-AzF cofactor $PrP^{Sc}$, as indicated.
